# Moisture Controls on Hydrogen Oxidizing Bacteria: Implications for the Global Soil Hydrogen Sink

**DOI:** 10.1101/2025.04.18.649571

**Authors:** Linta Reji, Matteo B. Bertagni, Fabien Paulot, Qianhui Qin, Xinning Zhang

## Abstract

Assessing the impact of increasing anthropogenic H_2_ emissions on Earth’s radiative balance depends on understanding the soil microbial H_2_ sink—the largest and the most uncertain term in the global H_2_ budget. A primary control regulating the soil sink is soil moisture, with a relationship that remains poorly constrained. Here, we assess the sensitivity of microbial H_2_ oxidation to soil moisture in laboratory experiments with three temperate soils—silty loam, sandy loam, and loamy sand. Using genome-resolved metagenomics, we link H_2_ oxidation dynamics in these soils to specific microbial taxa adapted to withstand desiccation that have differential contributions to H_2_ uptake along the moisture gradient. The experiments reveal a notably low moisture threshold for H_2_ oxidizer activity, at water potentials between –70 and –100 MPa across all soil types, including in an arid sandy soil. These measurements, which represent some of the lowest water potentials reported for soil microbial activity, point to atmospheric H_2_ as a vital resource for microbial survival under stressful conditions. Through global simulations, we further show that the low moisture threshold for microbial activation increases the contribution of arid and semi-arid regions for soil H_2_ uptake by 4-7pp, while decreasing the contribution of temperate and continental regions (−7pp), even when assuming a linear scaling between uptake potential and soil organic carbon, as suggested by our experiments. Our results highlight the importance of H_2_ uptake under extreme hydrological conditions, particularly the roles of desertification, dryland expansion, and H_2_-oxidizer ecophysiology in modulating long-term changes in H_2_ uptake.

## INTRODUCTION

Molecular hydrogen (H_2_) is an abundant trace gas in the troposphere, with a current global mean concentration of ~550 ppb in the marine boundary layer (1). H_2_ is expected to play a key role in the global clean energy transition due to its potential as a low-carbon fuel (2,3). However, an increasing reliance on H_2_ energy will likely raise atmospheric H_2_ concentrations, as H_2_ is leak-prone (4,5). This warrants a careful evaluation of the potential climate effects of increasing H_2_ levels in the atmosphere. While H_2_ is not a greenhouse gas, it still imparts a significant indirect radiative forcing due to its interaction with hydroxyl (OH) radicals in the atmosphere. This interaction prolongs the lifetime of methane in the troposphere and contributes to the production of stratospheric water vapor and tropospheric ozone (6–11). Given the significance of H_2_ for Earth’s radiative balance (100-year time-horizon Global Warming Potential, GWP^2^ 100, estimated to be 11.6 ± 2.8; (11)), the balance of the global H_2_ budget is a critical consideration in the context of accelerated energy transitions.

Uptake by soil microbes is the largest sink for atmospheric H_2_, accounting for over 70% of the global tropospheric H_2_ sink (12–14,8). By reducing the amount of H_2_ available to react with OH radicals in the atmosphere, the soil sink considerably dampens the indirect radiative impact of H_2_. Recent culture-based and genomic work has established that soil uptake is microbially mediated, with the high-affinity hydrogen oxidizing bacteria (HA-HOB) being widely distributed and active across ecosystems (15–21). The magnitude of the global microbial H_2_ sink is far from constrained and is the largest uncertainty in assessing the global warming potential of H_2_ (11).

One of the primary sources of uncertainty in soil sink estimates is the lack of constraint on the effects of soil moisture variability on H_2_ uptake (22–27). This is due to the complex ways in which shifts in soil moisture aeect H_2_ uptake, including direct impacts on microbial metabolism and the regulation of H_2_ diffusion into soil pores (24,26). At the dry end of the moisture spectrum, HA-HOB are water-stressed, resulting in limited H_2_ oxidation. Uptake is also low at higher moisture levels, due to the reduced diffusion of H_2_ in water-filled pores. Thus, the relationship between soil moisture and H_2_ uptake is highly non-linear (23,25,26).

Several attempts have been made to parameterize the soil sink to capture its sensitivity to soil moisture variability, along with other abiotic factors such as temperature and soil type (8,23,25,26,28,29). Due to the paucity of observational constraints on soil uptake (25,30–34), these parameterizations result in inaccurate estimates of the seasonality and spatial variability of uptake (8,27,35). Most parameterizations include a water-stress threshold, below which uptake is essentially zero; however, the magnitude of this threshold is poorly known (8,29). Similarly, an optimal moisture level at which peak uptake occurs has also been identified as a key threshold regulating microbial H_2_ oxidation, as above this level, uptake declines due to limited diffusion of H_2_ and O_2_ (26). How these moisture thresholds vary across soil types and HA-HOB respond to changes in moisture levels remain largely unknown. Furthermore, the unconstrained sensitivity of the H_2_ sink to soil moisture makes it challenging to capture spatio-temporal variability in sink strength in global models, especially with increasing atmospheric H_2_ levels (27).

In this study, we performed controlled laboratory experiments to decipher the mechanistic relationship between soil moisture and microbial H_2_ oxidation activity focusing on conditions of negligible diffusive limitation. We measured H_2_ oxidation rates in moisture controlled incubations of three soil types—a sandy loam, a silty loam, and a loamy sand. These soils were sampled from three distinct temperate ecosystems: a deciduous forest, a grass-shrub meadow, and an oak-pine forest, respectively. Based on the previously established non-linear relationship between soil moisture and H_2_ uptake, we expected the measured rates to be directly linked to the moisture response of high-affinity H_2_ oxidizers, particularly under drier conditions when uptake is not diffusion limited. Thus, in incubations set up to minimize diffusive limitation for uptake, we determined the moisture sensitivity of HA-HOB by coupling measurements of water potential and microbial H_2_ oxidation rates. We then revised previous formulations of soil H_2_ uptake models based on the experimental data, and assessed the response of the global soil H_2_ sink, both spatially and temporally, to the revised parameterization. We further analyzed the microbial community structure in our soil incubations to specifically examine how the diversity, abundance, and relative activity of different HA-hydrogenases (i.e., enzymes mediating high-affinity H_2_ oxidation) and HA-HOB vary with moisture changes and soil type.

## METHODS

### Soil collection, characterization, and experimental setup

Soils were collected from three distinct ecosystems: a deciduous forest, a managed grassland shrub meadow, and a sandy oak-pine forest. The forest and meadow ecosystems are located within the Watershed Research Institute in Pennington, New Jersey (40.35501 N 74.77458 W, and 40.35360 N 74.77433 W, respectively). The oak-pine forest is located within the New Jersey Pinelands/Pine Barrens, and we sampled in the Brendan T. Byrne State Forest (39.88683 N 74.53663 W). At each location, soil samples were collected from the top ~ 10 cm, after removing the organic layer, if present. Samples were collected in pre-cleaned (acid-washed and UV-sterilized) sealable plastic bags and stored on ice while transporting back to the laboratory. Once in the lab, soils were spread into a thin layer on a clean tray lined with an acid-washed and UV-sterilized plastic bag, any large plant material was manually removed, and the samples were left to air-dry for up to 3 days (until visually dry). The air-dried soils were sieved through a clean 2 mm sieve and stored in pre-cleaned sealable polyethylene bags in the refrigerator. All moisture incubations were set up within 10-15 days of sieving and storing the soil.

Before setting up the moisture incubations, the sieved soils were characterized for soil texture, pH, porosity, bulk density, and gravimetric moisture content. Soil texture analysis revealed the forest soil to be sandy loam, the meadow soil silty loam, and the Pine Barrens (herein PB) soil loamy sand. All three soils were acidic, with pH values between 4-4.6 (forest soil=4.2, meadow soil=4.6, and PB sand=4.1). Measured porosity values for the sieved soils ranged from ~0.6 for both the meadow and forest soils and ~0.4 for PB sand. Gravimetric moisture content (i.e., g water/g dry-soil) was determined by oven-drying the soil at 105 ^0^C until constant weight. Volumetric moisture content was then estimated by multiplying gravimetric moisture content by bulk density.

Soil organic matter content was estimated using the loss on ignition (LOI) method. Briefly, air-dried soil samples in triplicate were oven-dried at 105°C until constant weight to remove moisture. The dry weight was recorded, after which samples were combusted in a muffle furnace at 505°C for 4 hours to oxidize organic matter. LOI (%) was calculated as:

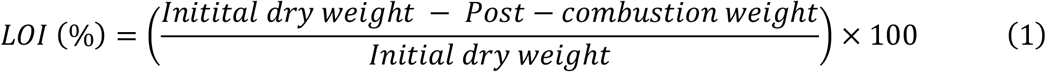

LOI values were converted to soil organic carbon as:

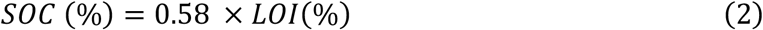

To set up the moisture-amended incubations, we added a small amount of air-dried soil (2 g each of forest and meadow soils, and 4 g of PB sand) into acid-washed and autoclaved 240 ml serum bottles. We then slowly added ultrapure water to the soil to achieve specific moisture levels (3 replicates per moisture level), mixed by gently shaking the bottle. A sterile autoclaved polypropelene spatula was occasionally used to gently mix the soil to facilitate water distribution. We then quickly sealed the bottle with parafilm to reduce evaporative losses, wrapped the bottle in aluminum foil, and incubated at 22 ^0^C. The acclimation period lasted an average of 6 days (i.e., all measurements were taken within 5-7 days of incubation). Any evaporative water loss during the incubation period was assessed by weighing the bottles daily. No measurable weight loss occurred during the incubation period. We further determined that the experiments were not diffusion-limited in the dry range (Fig. S1), as described in Supplementary Methods.

To extrapolate the observations to arid soils, we collected soils from two locations in Arizona (AZ soil 1: 34°44′12.12″ N, 111°59′7.488″ W; AZ soil 2: 33°58′9.528″ N, 112°7′45.251″ W). Texture for these soils were determined to be loamy sand and sandy loam, respectively. These soils were incubated in similar conditions (2g soil in 240 ml serum bottles), with adjusted moisture contents ranging from ~6-11% saturation, and were acclimated overnight before H_2_ oxidation measurements.

### Microbial H_2_ consumption measurements and soil moisture characterization

For measuring H_2_ oxidation rates, incubation bottles were retrieved from the incubator, flushed with room air, and immediately closed with a butyl rubber stopper, pre boiled in sodium hydroxide, and crimped with aluminum seal. Experiments were conducted at ambient H_2_ concentrations at 22 ^0^C. Time zero sample (2 mL) was immediately drawn from the bottle headspace using a luer-lock syringe and injected into a reducing compound photometer (RCP) gas chromatograph (Peak Laboratories LLC) fitted with a 1 mL sample loop. Additional headspace samples (2 mL) were collected and immediately measured on the GC-RCP until an exponentially decreasing trend in H_2_ concentrations was captured.

Triplicate incubations for each soil type-moisture combination were measured in parallel, with headspace samples sequentially drawn and analyzed from each replicate. Most experiments lasted under 2 hours. The AZ soils were much slower in consuming H_2_, and therefore, experiment duration was substantially longer for these soils (550 min for AZ soil 1 and 3000 min for AZ soil 2). The limit of detection for the measurements was determined to be 57 ppb. Based on this, the limits of detection for H_2_ oxidation rates were approximately 0.0001 ppm min^−1^ g-Soil^−1^ (for incubations containing 2g soil; forest and meadow soils), 0.00004 ppm min^−1^ g-Soil^−1^ (for PB soil incubations containing 4g soil), 0.00005 ppm min^−1^ g Soil^−1^ for AZ soil 1, and 0.00001 ppm min^−1^ g-Soil^−1^ for AZ soil 2. H_2_ drawdown in the headspace was modeled as a first-order decay process, and the rate constants (min^−1^) were used as proxy for the rate of H_2_ oxidation. For each soil type, killed controls (i.e., containing autoclaved airdried soil) and no-soil (empty) controls were also measured for H_2_ uptake (Fig. S2).

Immediately following uptake measurements, the bottles were briefly sealed with parafilm again and re-weighed to determine any evaporative losses during measurement. We did not observe any measurable weight loss during measurements. Soil was quickly subsampled into pre-weighted receptacles for: (i) re-measuring gravimetric water content, (ii) water potential measurements, and (iii) DNA/RNA. Approximately 1-2 ml of soil from the incubations was used for water potential measurements using the WP4C dewpoint potentiometer (METER Group).

### Nucleic acid extractions and sequencing

Total nucleic acids (DNA and RNA) were co-extracted from ~0.4 g of soil by combining the RNeasy PowerSoil Total RNA kit with the RNeasy PowerSoil DNA Elution kit (Qiagen), using manufacturer’s protocols. Quality and yield of the extracted DNA and RNA samples were assessed using NanoDrop and Qubit fluorometer respectively. For the forest soil, replicate soil samples from each moisture perturbation were extracted separately as these samples generally yielded high DNA/RNA concentrations. In contrast, for the Meadow soil and Pine Barrens sand, triplicate soil samples per moisture amendment were pooled prior to extraction. DNA extracts were cleaned up using the QIAGEN DNeasy PowerClean Pro Cleanup kit to improve quality, and were re-assessed on the NanoDrop to ensure that impurity levels were minimal in the final extracts. Extracted nucleic acids were stored at −80C until further processing.

DNA and RNA samples corresponding to 4-5 moisture levels for each soil type were used for metagenome and metatranscriptome sequencing (Table S1). Total DNA and RNA samples for the selected samples were sent to the Princeton Genomic Core facility for paired-end sequencing on the Illumina NovaSeq platform (S1 300nt flowcell). For the RNA samples, a ribo-depletion step was performed prior to library construction. The resulting reads were adapter-trimmed and demultiplexed by the sequencing facility. Further processing of the reads was carried out using the Princeton Research Computing cluster.

### Processing and analyses of metagenomes and metatranscriptomes

Demultiplexed metagenomic reads were quality filtered using FastQC (v0.11.9; (36)) and Trimmomatic (v0.39; (37)). Metagenomes for each soil type were co-assembled using MEGAHIT (--min-contig-len 1000; v1.2.9; (38,39)). Bowtie 2 (v2.3.5; (40)) was used to map reads back to the assembled contigs. Contig gene-calling was performed by using Prodigal (41) and the resulting gene sequences were used as input for DIAMOND blastp (v. 2.1.10; (42)) search against the UNIPROT-SWISSPROT database to obtain functional annotations. Contigs longer than 1500 bp were binned using MetaBAT 2 (v1.12.1; (43)) and MaxBin 2 (v2.2.7; (44)). Bin refinement was carried out using the bin_refinement module in MetaWRAP (v1.2; (45)). Prodigal (v2.6.3; (41)) was used to obtain amino acid sequences of gene calls for each bin. CheckM (v1.2.3; (46)) was used for bin quality assessment. We only retained medium-to high-quality (>70% complete and <10% contamination) genomes for downstream analyses. Taxonomic annotations were obtained by using the Genome Taxonomy Database toolkit (v.2.4.0; (47,48), classifying against the R220 database.

Metatranscriptome reads were QC-filtered to remove adapters by using FastQC (v0.11.9; (36)) and Cutadapt (v3.7; (49)). Using Bowtie 2 (v2.3.5; (40)), QC-filtered reads were mapped to the metagenomic contigs obtained for each soil type. Relative abundance estimates for contigs and metagenome-assembled genomes (MAGs) were obtained by using CoverM (v0.7.0; (50)). For abundance estimates of MAGs in the metagenomes, Coverm “-- method rpkm” was used. Conversely, “--method tpm” was used for estimating relative transcription of the MAGs in the metatranscriptomes. Additional functional annotations for MAGs were obtained via DRAM (v0.1.2; (41)) and RASTtk (v1.073; (52)) as implemented within the KBase platform (53). KO annotations for translated proteins in each MAG were obtained using GhostKOALA (54). DIAMOND BLASTp (42) searches against databases generated for each hydrogenase group from HydDB (55) were used identify putative hydrogenase homologs in MAGs and contigs. Hits were aligned with reference sequences using Mafft (v7.475; (56)), which were filtered using trimAl (v1.4.1; “-gt 0.8 -cons 60”; (57)), then used to compute phylogenetic trees using FastTreeMP (v2.1.10; (58)). Potential false positives were filtered out based on phylogenetic placement, homology, and residue comparisons.

### Global estimates of H_2_ deposition velocity, v_d_(H_2_)

The deposition velocity of H_2_ (v_d_(H_2_)) is calculated with the solution of the vertically explicit H_2_ mass balance in soil (59,60) as:

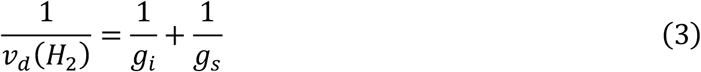

where *g*_*i*_ represents the H_2_ conductance through possible diffusive barriers (e.g., snow cover, canopy), which reduces H_2_ transport to HA-HOB, and g_s_ is the conductance through the active soil layer where H_2_ oxidation by HA-HOB occurs. v_d_(H_2_) in this model accounts for both biotic and physical processes regulating the soil sink.

g_s_ *g*_*s*_ an be expressed as:

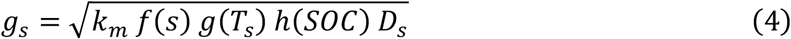

where f(s), g(Ts) (23), and h(SOC) are the sensitivity of microbial H_2_ oxidation to volumetric soil moisture (s), temperature (Ts), and SOC, respectively (26). D_s_ is the diffusion of H_2_ in the soil and k_m_ is the maximum oxidation rate of H_2_ by HA-HOB. f(s) is parameterized following Bertagni (2021; (26)) as:

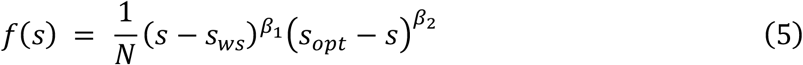

where s_ws_ is the lower moisture threshold for bacterial activity, and s_opt_ is the optimum. These soil-specific moisture values can be derived using only two water potential values (for bacterial inhibition and optimum) and the soil water characteristic curves. For our simulations, these water potential values are derived from the experiments. N is a normalization factor such that max(f)=1. *h*(*SOC*) was empirically derived based on the experiments performed in this study as a linear fit between maximum of H_2_ oxidation rate for each soil type and mean SOC (%) (described in Results).

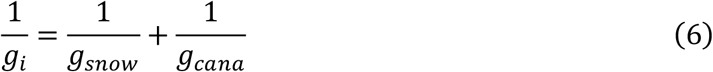

g_i_ reflects the transport of H_2_ through the canopy (g_cano_) and through snow (g_snow_). g_snow_ is calculated from snow depth assuming a snow porosity of 0.645. g_cano_ is prescribed following (61). Given the importance of short-term hydrological fluctuations (26), we use hourly soil moisture, temperature, and pressure from ERA5 reanalysis (62,63). The time-invariant spatial distribution of SOC is prescribed following Soil Grids v2.0 (top 5cm, (64)).

*k*_*m*_ is optimized for each configuration, such that the global deposition average v_d_(H_2_) is 0.09 mm/s for year 2010 (8).

All statistical analyses and plotting were done in R (v4.4.1; (65)) and Python. Plots were saved as PDF files, and final figures were generated after minor editing and combining of the plots in InkScape.

## RESULTS

### Moisture effects on microbial H_2_ oxidation across soil types

Here we report microbial H_2_ consumption measurements under controlled moisture conditions in the three soil types—forest sandy loam, meadow silty loam, and PB loamy sand. Hereafter, these are also referred to as forest, meadow, and PB soils, respectively. Following previous literature, we first report the results in terms of soil moisture; we then show that using the soil water potential rather than soil moisture offers better quantitative insights on the shared water-stress limitations for HA-HOB across soil types. Specifically for soil moisture, here we express the experimental saturation as the fraction of water-filled soil pores, where 100% represents saturation and 0% corresponds to fully dried (oven-dried) conditions. However, since even after oven-drying, residual water remains trapped in smaller soil pores, we stress that 0% does not signify a complete absence of water – a known problem in the soil hydrology community (66). As later discussed, this is an important aspect to consider, especially when evaluating the moisture percentages in the dry range.

Experiments revealed the expected non-linear relationship between soil moisture and H_2_ oxidation activity in all three soils, with maximum uptake occurring below 40% saturation (Fig. 1a). H_2_ consumption was not observed in control incubations (i.e., autoclaved soils, air-dried soils, and incubations without soils; Fig. S1). Measurable H_2_ oxidation was observed under very low moisture levels, only slightly above air-dried conditions (Fig. 1a). Oxidation rates initially increased sharply with increasing moisture until reaching a maximum (Fig. 1a). Peak activity was observed at ~20%, 25% and 35% saturation in the PB, meadow and forest soils, respectively. Rates decreased above these moisture levels, indicating reduced microbial consumption of H_2_ (Fig. 1a), and slight diffusive limitation above 60% saturation (Fig. S1). In PB sand, a clear decrease in uptake rate was not evident, as higher targeted saturation levels resulted in similar uptake rates and, as later shown, in soil water potential. This pattern indicates drainage from the sandy soil, as expected at saturations above 30%, impeding higher saturation levels.

**Figure 1:**
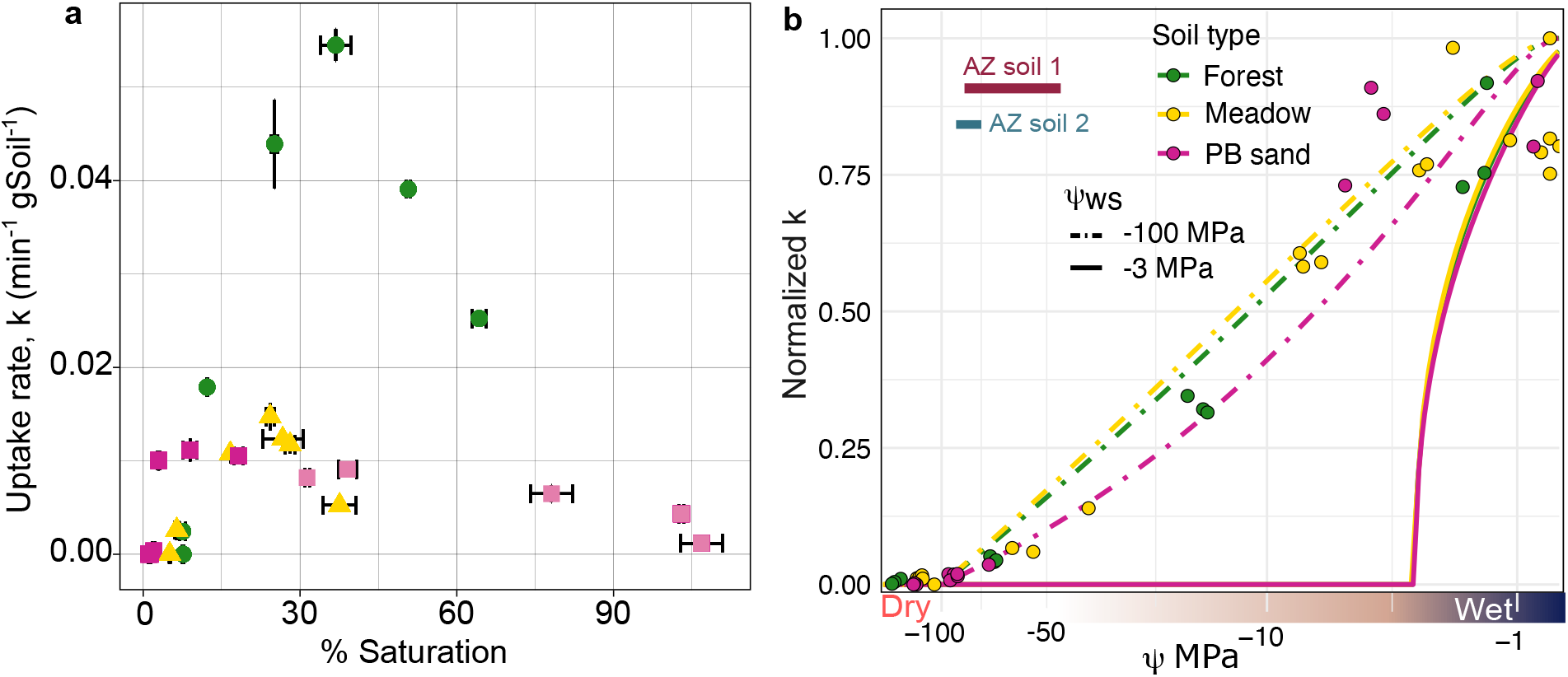
Relationship between soil moisture and microbial H_2_ oxidation. **(a)** Average H_2_ oxidation rates (min^−1^ g-soil^−1^) of triplicate incubations versus mean saturation (%). Saturation levels were estimated as volumetric moisture content/porosity. Error bars are standard errors around the mean. Faded fill colors for PB sand indicate that the incubations did not show slower H_2_ oxidation with rising moisture >~30% saturation, likely owing to a dieerence between targeted and real saturation in the experimental setup. (**b**) Average uptake rates (min^−1^) normalized by maximum measured uptake rate in each soil type versus measured water potential Ψ(MPa) of triplicate incubations. The curves are modeled uptake rates *f*(*S*) ∗ *g* (*T*_*S*_), based on parameterizations developed in (26), revising the water-stress threshold (26) from −3 MPa to −100 MPa. The two shaded rectangles in (**b**) indicate the range between the highest measured Ψ with no H_2_ oxidation and the lowest Ψ at which H_2_ oxidation was observed for the two AZ soils (Table S2).

Across all three soils, the forest sandy loam soils showed the fastest microbial uptake (per gram soil) with the highest measured rate constant (min^−1^ g-soil^−1^) 4- and 5-fold higher than that of the meadow silty loam and PB loamy sand soils, respectively. Uptake was not observed under fully air-dried conditions, which were estimated to correspond to approximately 5%, 7.6%, and 1.3% saturation for meadow, forest, and PB sand soils, respectively (Fig. S2; Table 1). Measurable uptake was observed under moisture levels only slightly above air-dried conditions: ~6.5% saturation in silty loam (meadow), ~7.6% saturation in sandy loam (forest), and ~2% in PB sand (Table 1). However, there is large uncertainty in these moisture measurements in the very dry range due to the little amount of water present. For example, for the forest soil, the lowest moisture amendment for which uptake was detected was estimated to have the same moisture content as the air-dried soil at ~7.6% saturation (Table S1). This highlights methodological limitations in estimating moisture content by conventional methods (i.e., oven-drying the soil sample until constant weight). Residual water that may still remain in the soil makes estimating moisture content in the dry end of the spectrum especially challenging. We therefore also measured the water potential (Ψ) of these soils as a more intrinsic measure of water availability.

### Unexpectedly low water stress threshold for high-affinity H_2_ oxidizers promotes H_2_ uptake in dry soils

Water potential (Ψ), a measure of the energy state of water, is widely used as an indicator of soil water availability to plants and microbes (e.g., (67–69)). Our water potential measurements indeed showed that the air-dried samples had much lower water availability than those amended with moisture, even in the very dry range (Table S1). Moreover, the measurements revealed a low and consistent moisture condition for the onset of microbial H_2_ consumption, which we define here as the water-stress threshold, Ψ_ws_. In the forest soil, measurable H_2_ oxidation occurred at Ψ ~ −70 MPa, with no uptake in the air-dried soils at −130 to −140 MPa (Table S1). A similar Ψ_ws_ was observed for the meadow soil, as the driest sample for which H_2_ oxidation was observed was at −65 MPa (Table S1). The PB sand soils were active under lower water potentials, down to approximately −90 MPa (Table S1). We performed a limited number of tests on two sandy soils from arid regions and these desert soils exhibited significantly lower H_2_ oxidation rates than the temperate soils. Notably, their Ψ_ws_ was similar to the other soils, as non-zero oxidation rates were measured when water potential exceeded −75 MPa (Fig. 1b, Table S2). Collectively, these results suggest the water stress threshold of HA-HOB to be between –70 and –100 MPa, among the lowest thresholds ever reported for microbial activity (67,68,70).

These measurements further allowed direct integration of our results into the recently developed mechanistic model for H_2_ uptake that leverages Ψ as a fundamental measure of moisture content (22; Fig. 1b). Notably, this approach eliminates the need for soil-specific moisture thresholds for water stress and optimal conditions in parameterizing H_2_ uptake as it directly links bacterial activity to thermodynamic condiserations on water availability. Revising Ψ_ws_ from −3 MPa, a value based on plant physiology used in the original model formulation (26), to −100 MPa measured here yielded significantly improved fits to the experimental data (Fig. 1b).

### Uptake rates increase with soil carbon content

For the two loamy soils (forest and meadow), H_2_ oxidation rates scaled with average soil organic carbon (SOC) content within the drier range of the moisture spectrum (Fig. S4a). Peak oxidation rate in the forest soil was approximately 4-fold higher than that in the meadow soil on a per gram basis, when not controlled for SOC content. However, this difference disappeared when the rates were scaled by mean SOC (g/g), as the scaled rates were essentially equivalent for the two loam soils below the optimal moisture thresholds (i.e., <30% saturation). The loamy sand (PB soil), in contrast, showed higher rates than the two loams when adjusted for SOC content (Fig. S4a), suggesting higher relative abundance and/or activity of HA-HOB in these soils. Maximum H_2_ oxidation rate (k_max_ min^−1^ gSoil^−1^) increased nearly linearly with SOC (Fig. S4b). A linear fit given by k_max_ = 0.00387 + 0.0024 * (SOC) (Fig. S4b) was used to incorporate a modulation by SOC to global simulations of v_d_(H_2_), as described next.

### Implications for the spatial distribution of the H_2_ sink strength and its historical change

To examine the implications of the observed moisture-sensitivity of microbial H_2_ oxidation rates on global H_2_ soil sink, we estimate v_d_(H_2_) (eq. 3) using a Ψ_ws_ of −3 MPa and −100 MPa. In this model, v_d_(H_2_) accounts for both biotic and physical processes regulating the soil sink. Simulations were run with and without modulation by SOC (Fig. S4b). We find that the particular value of the water stress threshold chosen for modeling significantly affects the spatial distribution of modeled uptake rates (Fig. 2a-c)(71). Soil moisture is predicted to be insufficient to support H_2_ uptake in ~90% of deserts and 45% of semi-arid regions for a Ψ_ws_ of −3 MPa. The degree of inhibition is reduced using our experimentally determined threshold of −100 MPa to 70 and 20%, respectively. In humid areas where moisture is generally high enough to support microbial H_2_ oxidation, v_d_(H_2_) is primarily controlled by diffusion (26). A lower Ψ_ws_ for HA-HOB tends to shift H_2_ uptake from continental regions (−7 pp) to desert (+7 pp) and semi-arid (+4 pp) regions. The relative importance of these regions also impacts the predicted change in v_d_(H_2_) over the second half of the 20^th^ century (Fig. 2d). In continental and polar regions, the increase in v_d_(H_2_) is insensitive to Ψ_ws_, and reflects increasing H_2_ soil diffusion due to lower soil moisture, warming, and reduced snow cover (8,27). In contrast, drying in desert and semi-arid regions has increased the areal extent of regions where soil moisture is too low to support H_2_ uptake. As deserts play a larger role with a lower Ψ_ws_, this contrasting response results in a lower increase in v_d_(H_2_) (+~7%) with a low Ψ_ws_ of −100 MPa, compared to a high Ψ_ws_of −3MPa (+~10.5%) from 1950 to 2019.

**Fig. 2.**
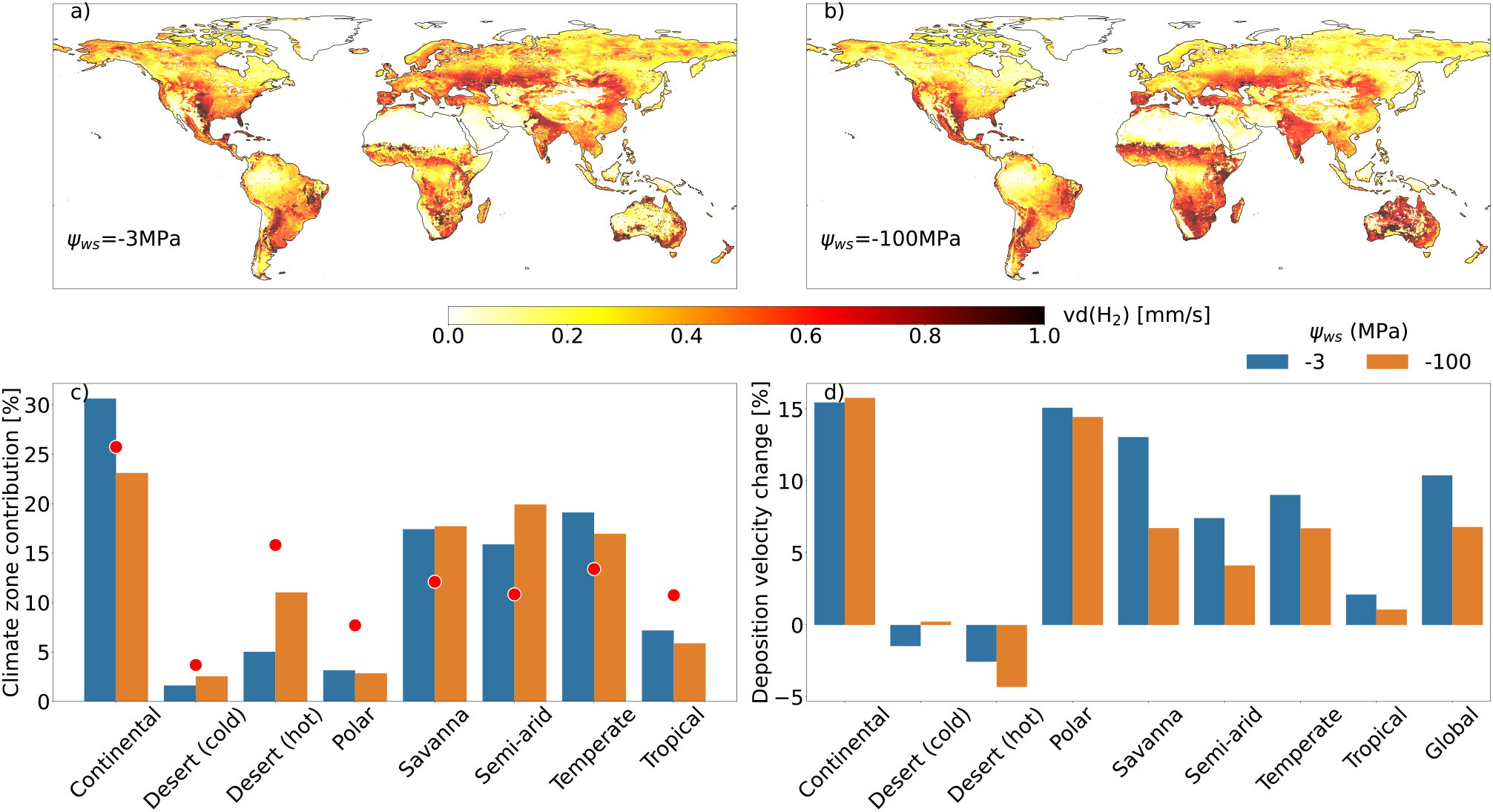
Impact of a lower moisture threshold for activation on global H_2_ uptake. Spatial distribution of H_2_ deposition velocity (v_d_(H_2_) mm/s) for a water stress threshold of (**a**) −3 MPa vs (**b)** −100 MPa averaged from 2005 to 2019. Simulations without the SOC modulation are presented in Fig. S5. (**c)** Fractional contribution of different climate zones to v_d_(H_2_) for different water stress threshold. Red dots indicate the fractional landmass corresponding to each climate zone north of 60S. (**d)** Relative change in v_d_(H_2_) across different climate zones from 1950-1964 to 2005-2019.

The simulated contribution of hot deserts to the H_2_ sink strength is sensitive to SOC (Fig. 2; S5). Without SOC modulation, the contribution of these regions to the overall H_2_ sink increases by 75% and reduces the simulated increase in H_2_ sink by ~20% (from 1950 to 2019).

### High-affinity hydrogen oxidizers are abundant and differentially active across the moisture range

For each soil type, we obtained metagenomes and metatranscriptomes at several moisture levels across the tested range (Table S3) to assess the community dynamics and ecophysiologies of high-affinity hydrogen oxidizers. Putative high-affinity hydrogenase (HA Hase) homologs were identified in both the contigs and metagenome-assembled genomes (MAGs) (see Methods).

Across all three soils, group 1h hydrogenase (Hhy) was the most dominant form of HA-Hases; (15); Table S3). Several Group 2a hydrogenases (with potential high-affinity activity, (17,72)) were also recovered from the forest soil metagenomes, but these were significantly less abundant than the 1h hydrogenases (6 homologs of group 2a versus 317 of group 1h identified in the forest soil contigs; Table S3). To estimate the relative abundance of HA-HOB in these soils, we compared the normalized abundances of HA-Hases with average normalized abundances of the single-copy marker genes for DNA gyrase, *gyrA* and *gyrB*. This analysis indicated that approximately 34% of the community in forest soil, ~25% in meadow soil, and 40% in PB sand have the ability for atmospheric H_2_ oxidation (Fig. S6). These estimates are comparable to previous surveys of HA-HOB in various ecosystems, which indicated the relative abundance of hydrogen oxidizers to be range from 20 to 57% of the total community across diverse ecosystems (73,74).

### Differential contribution of HA-HOB lineages to H_2_ oxidation across the moisture range

High-quality MAGs encoding HA-Hases (all group 1h) from the three soils were classified within the phyla *Acidobacterota, Actinomycetota, Eremiobacterota*, and *Chloroflexota* (Fig. 3a). Despite recent evidence indicates atmospheric H_2_ oxidation by Archaea (75), we did not find any archaeal MAGs with HA-Hase homologs. The specific genera within each phylum encoding HA-Hases were different across soil types (Fig. 3a). Relative abundances of several genera showed notable variations with moisture change. For example, in the forest soil, the *Acidobacterota* genus UBA5704 was more abundant than all other HA-HOB genera under drier conditions (Fig. 3a). This MAG is also the most highly transcribed among all HA-HOB MAGs in the forest soil, and its relative transcription increased with soil moisture (Fig. 3a). The *Actinomycetota* and *Eremiobacterota* genera (Bog-473 and DAIDGS01 respectively) showed increasing relative abundances in both the metagenomes and metatranscriptomes with increasing soil moisture (Fig. 3a). Several different lineages of *Acidobacterota* in the PB soil encode group 1h hydrogenase (Fig. 3a). Their relative abundance patterns suggest that different lineages mediate H_2_ uptake across varying moisture levels—in the driest sample sequenced, metatranscriptome read recruitment revealed transcriptional activity in only one of the four acidobacterial genera (Bog-105; Fig. 3a). In the meadow soil, uptake activity under low moisture conditions appears to be mediated by a novel genus within phylum *Actinomycetota* (Fig. 3a).

**Figure 3:**
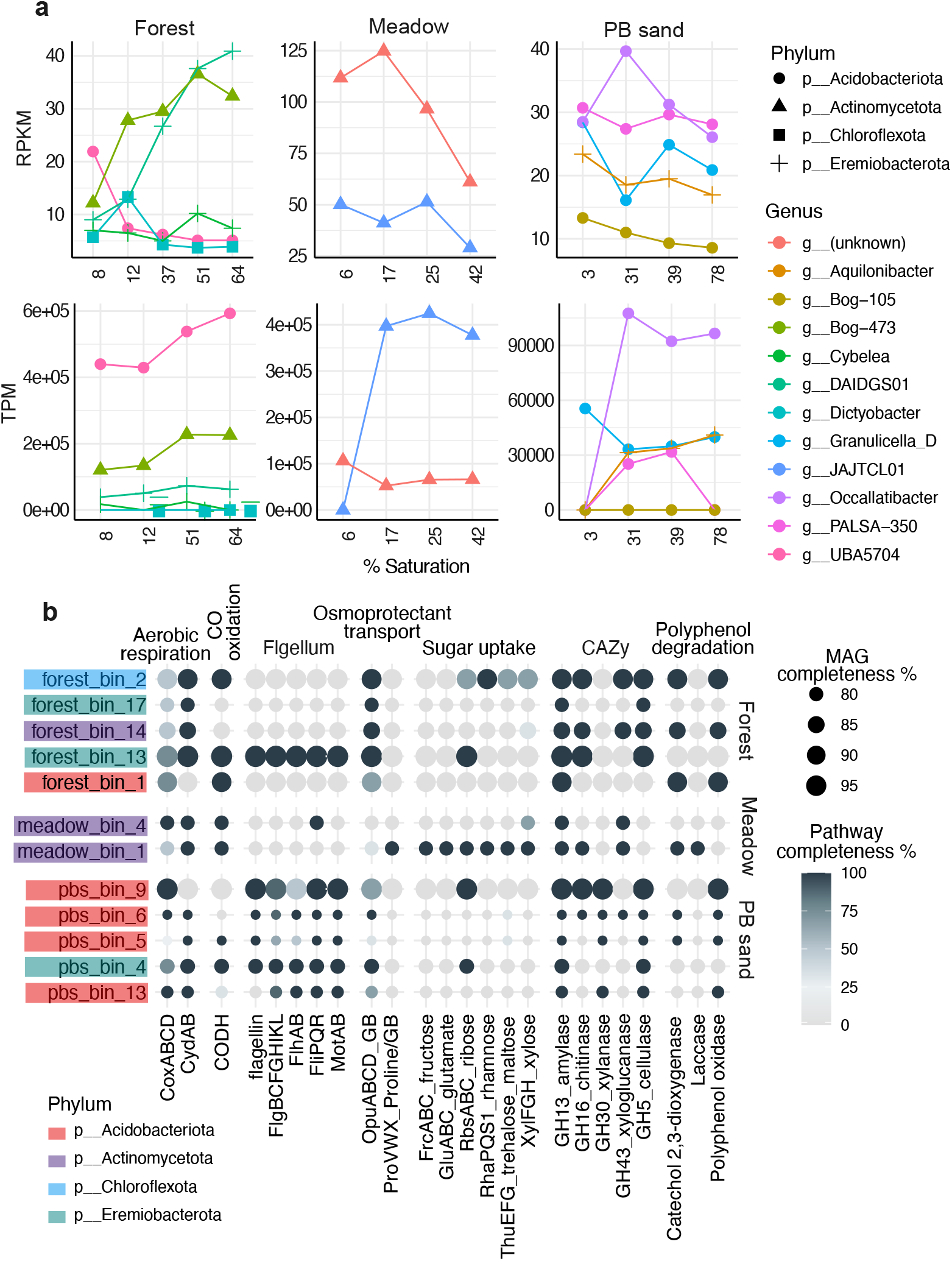
**a** Relative abundances and transcription (top and bottom panels, respectively) of HA-hydrogenase-encoding MAGs across moisture levels in forest soil, meadow soil, and PB sand. “g__” indicates a genus-level taxonomic identity. Relative abundances in the metagenomes are expressed as number of reads mapped per kilobase of genome per megabase of metagenome (RPKM). Relative transcriptional activity is expressed as the number of reads mapped per million total reads (transcripts per million, TPM). b Key metabolic pathways identified in the MAGs. Pathway completeness is expressed as proportion of identified gene subunits corresponding to each pathway, and the subunits considered are indicated in the figure. CO: carbon monoxide, GB: glycine betaine.

Functional profiling of the recovered HA-HOB MAGs revealed that aerobic heterotrophy was the predominant metabolic trait (Fig. 3b). While the capacity for the uptake of simple sugar compounds is relatively sparsely distributed, many of the MAGs harbor the potential for the degradation of complex carbon, including polyphenols and polysaccharides such as starch, chitin, xylan, and cellulose (Fig. 3b). These observations align with previous findings identifying HA-HOB as key players in the breakdown of complex plant-derived carbon in soils (76,77). Flagellar motility is widespread among the PB sand MAGs (Fig. 3b)—likely advantageous in the highly varying moisture regimes characteristic of sandy soils with low water retention. Additionally, nearly all MAGs recovered from the PB sand and forest metagenomes also possess the ability for osmoprotectant transport, specifically for glycine betaine (Fig. 3b), which likely enables these HA-HOB to resist desiccation under dry conditions. H_2_ oxidation likely provides an additional way to resist desiccation via the production of metabolic water, as has been hypothesized before (78,73). Notably, many of the MAGs also possess the ability for carbon monoxide oxidation (Fig. 3b), highlighting their capacity for versatile trace gas metabolism.

## DISCUSSION

Most parameterizations of the H_2_ soil sink include a water-stress or activation threshold for HA-HOB activity, below which there is no H_2_ uptake. To account for the variability in moisture content with soil texture, Bertagni et al., (2011; (26)) suggested the use of water potential as a more accurate representation of the actual water availability for hydrogen oxidizers in soils. A value of −3 MPa was recommended as the water-stress threshold as this is a typical value for the plant wilting point in semi-arid regions (26,79).

Following the discovery that HA-HOB are widespread in arid environments such as Antarctic desert soils (19), recent model implementations have used an even lower threshold of −10 MPa (27). The specific parameterization of the moisture threshold substantially affects model outputs, which makes it challenging to extrapolate spatial and temporal variability in uptake and attribute recent increases in atmospheric H_2_ levels to specific drivers (27).

Our results indicate a substantially lower moisture threshold for microbial H_2_ oxidation (closer to −100 MPa; Table 1 and Figure 1). These are some of the lowest water potential values reported for microbial activity in soils, comparable only to previous observations of CO oxidation in saline volcanic ash soils (down to −117 MPa, (68,70). Moisture thresholds have been proposed and experimentally determined for other microbial processes such as respiration and nitrification, but these values are generally much higher. For example, previous studies have identified −14 MPa as the threshold below which microbial respiration ceases (80,81). For nitrification, this threshold has been identified as −6 MPa (82). While the widespread occurrence of HA-HOB in hyper-arid cold (e.g., Antarctic) and hot (e.g., Atacama) deserts (19,78) points to a low water stress threshold for these microbes, our results represent the first time the moisture limits on HA-HOB activity have been quantified. The threshold values determined here are thus critical for developing a mechanistic understanding of the ecophysiology of HA-HOB, particularly in the face of global change. In addition, the observed similarity between the water-stress thresholds for uptake across all five soil types (forest sandy loam, meadow silty loam, PB loamy sand, and two arid soils—a loamy sand and sandy loam desert sands; Tables S1 and S2) is promising for global extrapolation of these results when modeling the soil H_2_ sink. With a lower threshold for activation, the importance of arid regions in H_2_ uptake increases significantly, which is noteworthy given the projected expansion of drylands with climate change (83,84).

Other key aspects of current global model parameterizations include the assumptions that HA-HOB are equally distributed across soil types(8). Similar to the results of numerous previous microbial surveys in diverse ecosystems, our microbial analyses suggest that this assumption is a gross oversimplification (Fig. 3). The relative abundances and activity of HA-HOB vary with soil and ecosystem type, aridity gradients, and SOC content (Fig. 3, S6; e.g., (18,19,74,78,85–87)). Thus integrating a more accurate representation of their population dynamics and ecophysiologies in global H_2_ models may help explain the difference in predicted activity between deserts versus temperate ecosystems (89,90).

Our results also show a positive correlation between SOC and H_2_ activity (Fig. S4), aligning with previous observations (56). Integrating this SOC scaling—or even more complex parameterizations such as that proposed by Karbin et al. (89)—could further reduce uncertainties in soil H_2_ oxidation potential. However, the overall impact of this modulation on H_2_ deposition velocity (the flux as seen from the atmosphere) remains relatively minor compared to the dominant influence of soil moisture (Fig. 2 and Fig. S5), which remains the primary driver of soil H_2_ uptake dynamics due to physical (i.e., diffusion) and biotic (i.e., bacterial activity) impacts.

From a microbial point of view, carbon-rich soils (e.g., forest soil) may exhibit more intense resource competition, leading to higher reliance of the microbial community on alternative energy sources such as H_2_. A majority of the forest soil HA-HOB MAGs also appear to be specializing in complex carbon degradation, suggesting their occupance of an oligotrophic niche (Fig. 3b). Within an ecosystem, local gradients in SOC may select for oligotrophic H_2_ oxidizers, as seen in desert soils (87). In general, the specific reasons for microbial H_2_ oxidation remains unclear even though various lines of evidence point to this being a survival strategy under resource limitation (73). On the drier end of the moisture spectrum, it has been proposed that H_2_ oxidation may provide metabolic water that could enhance survival (13). Yet water addition has also been demonstrated to substantially stimulate H_2_ oxidation activity in desert soils (19), highlighting the role of H_2_ as an energy source for microbes in oligotrophic systems. Our genome-centric analyses reveal that HA HOB are likely able to withstand desiccation by using osmoprotectants (Fig. 3b), further explaining their activity in extremely dry soils. Further studies are required to decipher the complex mechanisms regulating the ecophysiological response of HA-HOB to moisture variability.

Soil moisture and its sensitivity to environmental changes remain challenging to represent in global models (91), producing conflicting projections regarding future changes in aridity (92). With a lower moisture threshold that expands the importance of arid lands for H_2_ uptake, reducing uncertainty in future aridity changes will be important for global v_d_(H_2_) projections. In this regard, another important variable to consider is SOC changes. Warming is expected to reduce average soil carbon globally (93,94)—any effect this will have on v_d_(H_2_) remains unclear due to unknown mechanisms governing HA-HOB activity and SOC variability, both in abundance and form. Incorporating a linear SOC modulation increases the role of continental regions, which are not moisture-limited, in H_2_ uptake by nearly 35% (Fig. 2). Better constraining the mechanistic links between SOC and HA-HOB distribution and activity will be crucial to improving predictions of spatial distribution of the soil H_2_ sink strength under global change. Such knowledge is also critical to advance historical reconstructions of atmospheric H_2_ (95), clarifying long-term trends and variations in the global H_2_ budget.

## DATA AVAILABILITY STATEMENT

All data and code files can be found in the GitHub repository: https://github.com/Linta-Reji/Reji2025_H2_SoilMoisture. Measured H_2_ oxidation rates and soil properties have also been uploaded to FigShare. (Please note that since the FigShare DOI will not be available until after manuscript submission, we are providing this dataset as Additional Review Material along with the manuscript files). Metagenomes and metatranscriptomes generated in this study will be deposited in the NCBI Sequence Read Archive (available upon publication).

## ACKNOWLEDGMENTS

This research has been supported in part by the Water Grand Challenge Award from the High Meadows Environmental Institute at Princeton University, the Carbon Mitigation Initiative (CMI) at Princeton University, and by the US Department of Energy, Office of Energy Efficiency and Renewable Energy (EERE), specifically the Hydrogen and Fuel Cell Technologies Office (award NA23OAR4310138). The views expressed herein do not necessarily represent the views of NOAA, the U.S. Department of Energy, or the United States Government. The work reported on in this paper was substantially performed using the Princeton Research Computing resources at Princeton University which is consortium of groups led by the Princeton Institute for Computational Science and Engineering (PICSciE) and Office of Information Technology’s Research Computing. This work was supported by the Office of Science, Office of Biological and Environmental Research, of the US Department of Energy under Award Numbers DE-AC02-05CH11231, DE-AC02-06CH11357, DE-AC05-00OR22725, and DE-AC02-98CH10886, as part of the DOE Systems Biology Knowledgebase.

